# Rapid metagenomic next-generation sequencing during an investigation of hospital-acquired human parainfluenza virus 3 infections

**DOI:** 10.1101/074500

**Authors:** Alexander L. Greninger, Danielle M Zerr, Xuan Qin, Amanda L. Adler, Janet A. Englund, Keith R. Jerome

## Abstract

Metagenomic next-generation sequencing (mNGS) is increasingly used for the unbiased detection of viruses, bacteria, fungi, and eukaryotic parasites in clinical samples. Whole genome sequencing (WGS) of clinical bacterial isolates has been shown to inform hospital infection prevention practices, but the use of this technology during potential respiratory virus outbreaks has not been taken advantage of. Here, we report on the use of mNGS to inform the real-time infection prevention response to a cluster of hospital-acquired human parainfluenza 3 virus (HPIV3) infections at a children’s hospital. Isolates from 3 patients with hospital-acquired HPIV3 identified over a 12-day period on a general medical unit and 10 temporally-associated isolates from patients with community-acquired of HPIV3 were analyzed. Our sample-to-sequencer time was <24 hours while our sample-to-answer turn-around time was <60 hours with a hands-on time of approximately 6 hours. Eight (2 case isolates and 6 control isolates) of 13 samples had sufficient sequencing coverage to yield whole genomes for HPIV3, while 10 (2 cases and 8 controls) of 13 samples gave partial genomes and all 13 samples had >1 read to HPIV3. Phylogenetic clustering revealed the presence of identical HPIV3 genomic sequence in the two of the cases with hospital-acquired infection, consistent with the concern for recent transmission within the medical unit. Adequate sequence coverage was not recovered for the third case. This work demonstrates the promise of mNGS to provide actionable information for infection control in addition to microbial detection.

## Introduction

Healthcare-associated infections (HAIs) affect hundreds of millions of patients around the world and cost hospitals tens of billions of dollars each year (1). Hospital-acquired viral infections are a particular problem in pediatric settings, with upwards of one-third of HAIs attributable to viruses (2). Hospital acquisition of HPIVs have been recognized as a particular problem among patients with cancer, including pediatric patients (3). In one study, 80% of HPIV infections in pediatric cancer patients were acquired in the hospital (4).

Recently, next generation sequencing of pathogens from clinical samples has been used to inform infection prevention, identify antimicrobial resistance, and broaden differential diagnosis (5–10). WGS of clinical isolates has been most commonly used for genomic epidemiology of bacteria in hospital-acquired infections, where culture isolates are readily attainable, part of the normal workflow, and undergo minimal mutation during the culture process (11–13). WGS for viral pathogens is considered more difficult given the lack of routine culture and propensity of viral culture to induce mutations in viral genomes (7, 14). We used mNGS to rapidly sequence whole genomes from a putative cluster of HPIV3 to determine whether there was a common source on a medical unit. This is the first reported use of metagenomic next generation sequencing in real–time to inform hospital infection prevention.

### Outbreak Description

In June 2016, routine surveillance revealed three contemporaneous cases of hospital-acquired HPIV3 infection on a general medical unit. The first case was an 18-month old male with chronic lung disease who developed increased work of breathing on Day 109 of hospitalization. The patient required increased oxygen support and admission to the pediatric intensive care unit due to worsening hypoxemia. The patient was managed with a short course albuterol and steroids, and he recovered. Three days later a second case was identified. Patient 2, an 8-month old male with chronic lung disease developed increased work of breathing on Day 98 of hospitalization. The patient was also managed with a short course of albuterol and steroids and recovered. Then, 12 days after the first case, the third case was identified. Patient 3, a 4-month old male with congenital immunodeficiency developed cough and rhinorrhea on Day 22 of hospitalization. The patient’s symptoms resolved spontaneously without any interventions. Nasal swabs from all 3 patients tested positive for HPIV3 by RT-PCR.

### Outbreak Investigation

In response to the cluster, the medical unit leadership and hospital Infection Prevention Department conducted an investigation to identify potential sources of infection. Information about patient room locations, nursing assignments, involved provider teams, support services received by the patient, shared equipment, and patient movement in the hospital (to radiology, etc.) was collected. In addition, family members of patients and healthcare workers were queried about recent illness. The investigation identified that a healthcare worker with an upper respiratory tract infection had provided care to two of the healthcare-associated cases (patients 1 and 2).

In response to the event, nursing and physician staff received notification about the infections and education about the transmission of HPIV3. They were also reminded to stay home when ill. This information was included in weekly newsletters and daily unit-based huddles.

Isolates obtained from the 3 putative hospital-acquired HPIV3 cases and 10 control patients were sent to the University of Washington Virology Laboratory for mNGS two days after recognition of the cluster (18 days after the first patient was diagnosed with HPIV3). The original isolate for the third outbreak patient was no longer available so a second sample taken 6 days later was used for sequencing.

## Materials and Methods

### Setting

Seattle Children’s Hospital is a 316-bed quaternary care pediatric facility located in Seattle, Washington. The medical unit involved is located on one floor and includes 32 beds. Patients housed on the unit are newborn to 21 years of age and have a wide variety of acute and chronic health issues. Respiratory viral testing is routinely performed on symptomatic patients using nasal swabs evaluated using FilmArray Assay (BioFire, Salt Lake City, UT).

### Ethical Concerns

The study protocol was approved by the Children’s Institutional Review Board. Consent from patients was deemed unnecessary as all data already existed and were made anonymous for purposes of this study.

### Cluster and control isolates

Cluster isolates were clinical samples positive for HPIV3 by PCR from a patient hospitalized on the general medical unit who met criteria for hospital-acquired HPIV3 (clinical symptoms that developed >6 days after admission). These criteria were based on the incubation period of HPIV (15).

For strain comparison, clinical microbiology records were used to identify a convenience sample of control isolates from inpatient and outpatient children testing positive for HPIV3 by RT-PCR obtained during the same month as the cluster from patients with community-acquired HPIV3.

### mNGS library generation and sequencing

500uL of swab-inoculated viral transport media was filtered through a 0.45um Ultrafree-MC spin filter (Millipore) and 200uL of filtrate was used as input for extraction in a ZR Viral RNA Kit (Zymo Research). Extracted RNA was treated with Turbo DNAse (LifeTech) and first and second-strand synthesis was performed using random hexamers SuperScript III (Life Tech) and Sequenase enzynes (Agilent) (16). cDNA was purified using DNA Clean and Concentrator-5 kit (Zymo) and used as input for Nextera XT library generation with 20 cycles of PCR amplification. Sequencing libraries were purified using 1.0X AMPure XP beads (Beckman Coulter), quantified on the Qubit 3.0 (LifeTech) and Bioanalyzer 2100 (Agilent) (17). Samples were mixed equimolarly to achieve approximately 2 million reads per sample and sequenced using 1x180bp run on a MiSeq desktop sequencer (Illumina).

Sequencing reads were adapter- and quality-trimmed using cutadapt and mapped to the NCBI reference genome for HPIV3 (NC_001796) using Geneious v9.1 software (18). Consensus sequences of regions with >1X coverage were called by majority-voting base with hand curation. Genome alignments were made using MAFFT and phylogenetic trees were created using MrBayes (19, 20). HPIV3 genomes are deposited in Genbank under accessions KX574704-574711.

## Results

### Case patients

The cluster of 3 male patients with hospital-acquired HPIV3 ranged in age from 4-18 months. Additionally, all case patients had underlying medical conditions requiring frequent or prolonged hospitalization. None of the case patients had recent travel.

### Control patients

The 10 patients with community-acquired HPIV3 identified to serve as controls tested positive for HPIV3 by PCR within 15 days of the first case patient. Control patients were a median age of 9 months (range 2 weeks to 7 years) and 60% were male. Additionally, 60% of the control patients were previously healthy. One of the control patients (patient 9) had recent travel from the Midwest. All other patients came from western Washington state.

### Metagenomic sequencing reveals outbreak strain

A total of 22,904,287 quality/adapter-trimmed reads were derived from the 13 samples with a median trimmed read count of 1,666,529 reads per sample. The number of HPIV3 reads in each sample ranged from 4 reads to 202,946 reads (median 5,823 reads, Table). A total of 8 of 13 samples (2 case isolates and 6 control isolates) had sufficient coverage to yield near whole genomes for phylogenetic analysis. Phylogenetic analysis revealed that the 8 samples came from 6 distinct lineages, with the two hospital-acquired strains sharing the exact same sequence across the entire genome and identical to none of the control strains (Figure). Extending the tree to include 10 partial genomes from all samples with >50 reads mapping to HPIV3 (2 case isolates and 8 control isolates) showed 9 strains (Supplemental Figure 1). The third sample from a hospital-acquired case had 4 reads, each of which had identical sequence to the genomes recovered from the 2 outbreak strains (Supplemental Figure 2).

**Figure.**
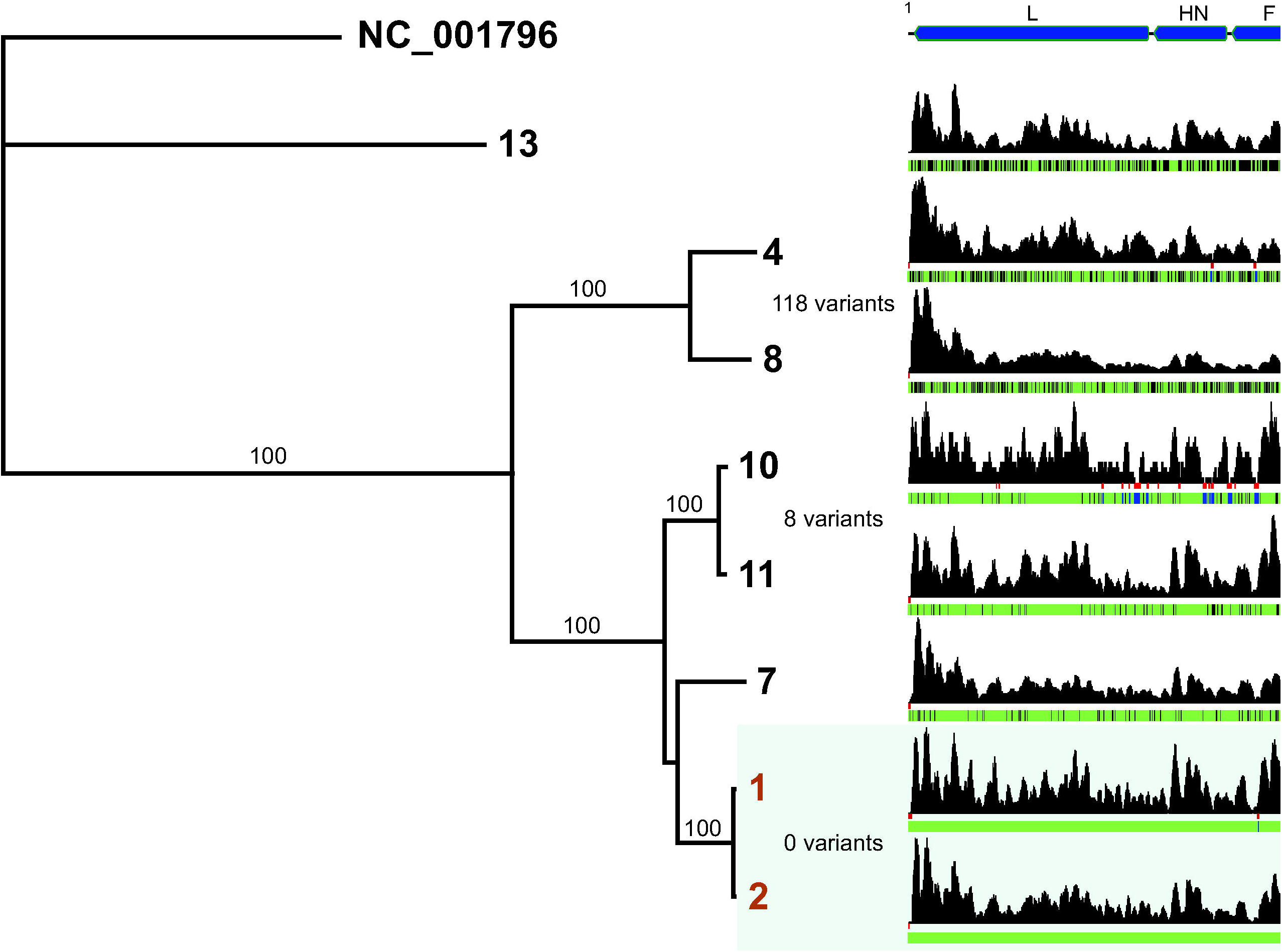
Whole genome phylogeny of HPIV3 sequences reveals identical sequence in putative outbreak samples. Near-complete genomes recovered from HPIV3 positive specimens were aligned by MAFFT and phylogenetic trees were constructed using MrBayes. Specimens are denoted by sample identifier name with the two identical clinical cases highlighted in orange. The NCBI reference genome for HPIV3 (NC_0001796) was used as an outgroup and its negative-stranded genome is highlighted in green. Consensus support values are denoted on branch labels. Normalized coverage maps are shown with no coverage areas shown in red and nucleotide changes relative to the outbreak sequence highlighted in black.

**Table.** Specimens sequenced in this study.

**Table: Specimens sequenced in this study**
| **Patient** | **Day positive**<br>**compared to**<br>**index case** | **Age** | **Sex** | **Filtered Reads** | **HPIV3 reads** |  |
| --- | --- | --- | --- | --- | --- | --- |
| 1 | 0 | 18 mo | Male | 1,814,866 | 5,823 | Hospital acquired |
| 2 | 3 | 8 mo | Male | 1,864,624 | 16,901 |  |
| 3 | 12 | 4 mo | Male | 1,097,618 | 4 |  |
| 4 | -8 | 2 yr | Male | 1,286,425 | 11,605 | Community acquired |
| 5 | -2 | 2 mo | Female | 1,960,840 | 21 |  |
| 6 | 1 | 1 yr | Male | 1,444,230 | 4 |  |
| 7 | 2 | 9 mo | Male | 1,565,065 | 132,401 |  |
| 8 | 3 | 2 wk | Female | 1,573,141 | 54,369 |  |
| 9 | 8 | 7 yr | Female | 1,054,320 | 88 |  |
| 10 | 10 | 22 mo | Female | 1,956,632 | 631 |  |
| 11 | 11 | 1 mo | Male | 1,666,529 | 31,502 |  |
| 12 | 13 | 2 mo | Male | 2,223,697 | 356 |  |
| 13 | 15 | 1 yr | Male | 3,396,300 | 202,946 |  |

BLASTN analysis of the outbreak strain revealed 100% nucleotide identity to HN genes from a 2011 Argentinian strain HNRG.036.11 (1146bp, KT765972), March 2010 Colombia strain FLI2812 (352bp, JQ268865), and 2012 Japan strain 70-PIV3 (147bp, AB831638). The closest whole genome sequence was from the October 2007 FLU8652 strain from Piura, Peru. BLASTN analysis of the farthest outgroup strain (patient 13) revealed 99.8% nucleotide identity to HN genes from the 1995 nosocomial outbreak strain from Massachusetts (427bp, AF039925) and 98.3% nucleotide identity to whole genome sequence from the HPIV3 cell culture strain 14702 originally isolated from Canada (15,462bp, EU424062).

Bayesian analysis has revealed an average 7.61×10^−4^ nucleotide substitutions per site per year across the HPIV1 genome (with 1.37×10^−3^ in the HN gene) (21). An analysis of HPIV3 strains in Japan showed approximately the same rate of evolution in the HN gene for HPIV3 (22). These data indicate approximately 11 sites change per year across the HPIV3 genome, suggesting the resolution of one nucleotide change in the HPIV3 genome in terms of transmission is approximately one month.

Here we report the first rapid use of metagenomics to inform infection prevention in a hospital setting. Prior approaches of using next-generation sequencing to inform infection control were entirely retrospective, focused on pure bacterial isolates, and/or utilized PCR tiling approaches based on a known organism (12, 13, 23–28). In this study mNGS retrieved identical whole genome HPIV3 sequences from two patients, confirming an association between the two cases and suggesting transmission from a single source. mNGS also recovered reads from a third outbreak patient that had identical sequence to the two patients in the cluster but the read coverage was not sufficient to determine with confidence that this case arose from the same source. Metagenomic sequencing was performed in a rapid turn-around time (<3 days) with approximately 6-8 hours of hands-on time. Linked bioinformatics analyses could potentially perform the same task from sample-to-answer in under 30 hours based on currently available Illumina sequencers and the depth of sequencing required. Nanopore sequencing could reduce this turnaround time even more; however, this method is not amenable to mNGS due to limited depth of sequencing (29, 23).

The main limitation of the approach taken here is the reduced sensitivity of mNGS relative to PCR (30). mNGS is not routinely able to recover whole genomes from FilmArray-positive specimens. In this study near complete genomes were recovered from only 8 of 13 specimens with a median of 1.7 million trimmed reads per sample, while partial genomes were recovered from 10 of 13 samples. One of the specimens for which whole genome HPIV3 sequence could not be retrieved was part of the putative cluster, reducing the ability to confidently link further transmission within the hospital. Of note, this specimen was obtained via a repeat swab approximately six days after the patient’s first positive test for HPIV3, and thus it is not surprising that little sequence could be recovered. In addition, whole genomes are not required to link cases as phylogenetic information can be recovered with only partial sequences (31).

A definitive advantage of mNGS is the use of only one protocol for detection and genome-wide analysis of a myriad of infectious diseases, which can aid in personnel training and reduction in turn-around time relative to other genome portioning methods. A potential limitation of this approach is cost, as we estimate that the marginal reagent cost of this sequencing was approximately $2000. However, the cost for mNGS versus capture sequencing methods is approximately the same and the temporal resolution and epidemiological information afforded by whole genome methods greatly exceeds that afforded by Sanger-based candidate gene sequencing.

Molecular tools enable the investigation of infection clusters and outbreaks, both in community and hospital settings. mNGS adds to the available tools and allows for unbiased pan-pathogen detection with single-nucleotide resolution that can enable detection and inference of transmission. In the hospital setting, this molecular data can play a critically important role in convincing healthcare workers and administrators that transmission is occurring and can provide rationale for expending resources and targeting interventions to prevent further transmission. The ability of metagenomic NGS to provide whole genome epidemiologic data in a rapid and actionable timeframe will advance clinical care as this technology moves from the research to the clinical setting.

---

**Supplemental Figure 1.** Partial genome phylogenetic analysis of samples with >50 HPIV3 reads reveals that additional community-acquired cases were not part of the hospital outbreak.

**Supplemental Figure 2.** Phylogenetic analysis of most informative 99bp trimmed read from patient 3 across all 10 samples for which sequence was available at that locus. Patient 3 clusters within the hospital outbreak; however, this limited sequence is not present in all low-coverage samples.

## References

1. 2011. Report on the Burden of Endemic Health Care-Associated Infection Worldwide. World Health Organization.

2. Aitken C, Jeffries DJ. 2001. Nosocomial spread of viral disease. Clin Microbiol Rev 14:528–546.

3. Shah DP, Shah PK, Azzi JM, Chemaly RF. 2016. Parainfluenza virus infections in hematopoietic cell transplant recipients and hematologic malignancy patients: A systematic review. Cancer Lett 370:358–364.

4. Maeng SH, Yoo HS, Choi S-H, Yoo KH, Kim Y-J, Sung KW, Lee NY, Koo HH. 2012. Impact of parainfluenza virus infection in pediatric cancer patients. Pediatr Blood Cancer 59:708–710.

5. Greninger AL, Messacar K, Dunnebacke T, Naccache SN, Federman S, Bouquet J, Mirsky D, Nomura Y, Yagi S, Glaser C, Vollmer M, Press CA, Kleinschmidt-DeMasters BK, Klenschmidt-DeMasters BK, Dominguez SR, Chiu CY. 2015. Clinical metagenomic identification of Balamuthia mandrillaris encephalitis and assembly of the draft genome: the continuing case for reference genome sequencing. Genome Med 7:113.

6. Naccache SN, Federman S, Veeraraghavan N, Zaharia M, Lee D, Samayoa E, Bouquet J, Greninger AL, Luk K-C, Enge B, Wadford DA, Messenger SL, Genrich GL, Pellegrino K, Grard G, Leroy E, Schneider BS, Fair JN, Martínez MA, Isa P, Crump JA, DeRisi JL, Sittler T, Hackett J, Miller S, Chiu CY. 2014. A cloud-compatible bioinformatics pipeline for ultrarapid pathogen identification from next-generation sequencing of clinical samples. Genome Res 24:1180–1192.

7. Greninger AL, Naccache SN, Messacar K, Clayton A, Yu G, Somasekar S, Federman S, Stryke D, Anderson C, Yagi S, Messenger S, Wadford D, Xia D, Watt JP, Van Haren K, Dominguez SR, Glaser C, Aldrovandi G, Chiu CY. 2015. A novel outbreak enterovirus D68 strain associated with acute flaccid myelitis cases in the USA (2012-14): a retrospective cohort study. Lancet Infect Dis 15:671–682.

8. Barzon L, Lavezzo E, Militello V, Toppo S, Palù G. 2011. Applications of next-generation sequencing technologies to diagnostic virology. Int J Mol Sci 12:7861–7884.

9. Capobianchi MR, Giombini E, Rozera G. 2013. Next-generation sequencing technology in clinical virology. Clin Microbiol Infect Off Publ Eur Soc Clin Microbiol Infect Dis 19:15–22.

10. Greninger AL, Runckel C, Chiu CY, Haggerty T, Parsonnet J, Ganem D, DeRisi JL. 2009. The complete genome of klassevirus — a novel picornavirus in pediatric stool. Virol J 6:82.

11. Gardy JL, Johnston JC, Ho Sui SJ, Cook VJ, Shah L, Brodkin E, Rempel S, Moore R, Zhao Y, Holt R, Varhol R, Birol I, Lem M, Sharma MK, Elwood K, Jones SJM, Brinkman FSL, Brunham RC, Tang P. 2011. Whole-genome sequencing and social-network analysis of a tuberculosis outbreak. N Engl J Med 364:730–739.

12. Snitkin ES, Zelazny AM, Thomas PJ, Stock F, NISC Comparative Sequencing Program Group, Henderson DK, Palmore TN, Segre JA. 2012. Tracking a hospital outbreak of carbapenem-resistant Klebsiella pneumoniae with whole-genome sequencing. Sci Transl Med 4:148ra116.

13. Roach DJ, Burton JN, Lee C, Stackhouse B, Butler-Wu SM, Cookson BT, Shendure J, Salipante SJ. 2015. A Year of Infection in the Intensive Care Unit: Prospective Whole Genome Sequencing of Bacterial Clinical Isolates Reveals Cryptic Transmissions and Novel Microbiota. PLoS Genet 11:e1005413.

14. Qiu S, Li P, Liu H, Wang Y, Liu N, Li C, Li S, Li M, Jiang Z, Sun H, Li Y, Xie J, Yang C, Wang J, Li H, Yi S, Wu Z, Jia L, Wang L, Hao R, Sun Y, Huang L, Ma H, Yuan Z, Song H. 2015. Whole-genome Sequencing for Tracing the Transmission Link between Two ARD Outbreaks Caused by a Novel HAdV Serotype 7 Variant, China. Sci Rep 5:13617.

15. Committee on Infectious Diseases. Red Book 2015.

16. Greninger AL, Chen EC, Sittler T, Scheinerman A, Roubinian N, Yu G, Kim E, Pillai DR, Guyard C, Mazzulli T, Isa P, Arias CF, Hackett J, Schochetman G, Miller S, Tang P, Chiu CY. 2010. A metagenomic analysis of pandemic influenza A (2009 H1N1) infection in patients from North America. PloS One 5:e13381.

17. Greninger AL, Chatterjee SS, Chan LC, Hamilton SM, Chambers HF, Chiu CY. 2016. Whole-Genome Sequencing of Methicillin-Resistant Staphylococcus aureus Resistant to Fifth-Generation Cephalosporins Reveals Potential Non-mecA Mechanisms of Resistance. PloS One 11:e0149541.

18. Martin M. 2011. Cutadapt removes adapter sequences from high-throughput sequencing reads. EMBnet.journal 17:pp. 10–12.

19. Katoh K, Misawa K, Kuma K, Miyata T. 2002. MAFFT: a novel method for rapid multiple sequence alignment based on fast Fourier transform. Nucleic Acids Res 30:3059–3066.

20. Huelsenbeck JP, Ronquist F. 2001. MRBAYES: Bayesian inference of phylogenetic trees. Bioinforma Oxf Engl 17:754–755.

21. Beck ET, He J, Nelson MI, Bose ME, Fan J, Kumar S, Henrickson KJ. 2012. Genome sequencing and phylogenetic analysis of 39 human parainfluenza virus type 1 strains isolated from 1997–2010. PloS One 7:e46048.

22. Mizuta K, Tsukagoshi H, Ikeda T, Aoki Y, Abiko C, Itagaki T, Nagano M, Noda M, Kimura H. 2014. Molecular evolution of the haemagglutinin-neuraminidase gene in human parainfluenza virus type 3 isolates from children with acute respiratory illness in Yamagata prefecture, Japan. J Med Microbiol 63:570–577.

23. Quick J, Loman NJ, Duraffour S, Simpson JT, Severi E, Cowley L, Bore JA, Koundouno R, Dudas G, Mikhail A, Ouédraogo N, Afrough B, Bah A, Baum JHJ, Becker-Ziaja B, Boettcher JP, Cabeza-Cabrerizo M, Camino-Sánchez Á, Carter LL, Doerrbecker J, Enkirch T, García-Dorival I, Hetzelt N, Hinzmann J, Holm T, Kafetzopoulou LE, Koropogui M, Kosgey A, Kuisma E, Logue CH, Mazzarelli A, Meisel S, Mertens M, Michel J, Ngabo D, Nitzsche K, Pallasch E, Patrono LV, Portmann J, Repits JG, Rickett NY, Sachse A, Singethan K, Vitoriano I, Yemanaberhan RL, Zekeng EG, Racine T, Bello A, Sall AA, Faye O, Faye O, Magassouba N, Williams CV, Amburgey V, Winona L, Davis E, Gerlach J, Washington F, Monteil V, Jourdain M, Bererd M, Camara A, Somlare H, Camara A, Gerard M, Bado G, Baillet B, Delaune D, Nebie KY, Diarra A, Savane Y, Pallawo RB, Gutierrez GJ, Milhano N, Roger I, Williams CJ, Yattara F, Lewandowski K, Taylor J, Rachwal P, Turner DJ, Pollakis G, Hiscox JA, Matthews DA, O’Shea MK, Johnston AM, Wilson D, Hutley E, Smit E, Di Caro A, Wölfel R, Stoecker K, Fleischmann E, Gabriel M, Weller SA, Koivogui L, Diallo B, Keïta S, Rambaut A, Formenty P, Günther S, Carroll MW. 2016. Real-time, portable genome sequencing for Ebola surveillance. Nature 530:228–232.

24. Gire SK, Goba A, Andersen KG, Sealfon RSG, Park DJ, Kanneh L, Jalloh S, Momoh M, Fullah M, Dudas G, Wohl S, Moses LM, Yozwiak NL, Winnicki S, Matranga CB, Malboeuf CM, Qu J, Gladden AD, Schaffner SF, Yang X, Jiang P-P, Nekoui M, Colubri A, Coomber MR, Fonnie M, Moigboi A, Gbakie M, Kamara FK, Tucker V, Konuwa E, Saffa S, Sellu J, Jalloh AA, Kovoma A, Koninga J, Mustapha I, Kargbo K, Foday M, Yillah M, Kanneh F, Robert W, Massally JLB, Chapman SB, Bochicchio J, Murphy C, Nusbaum C, Young S, Birren BW, Grant DS, Scheiffelin JS, Lander ES, Happi C, Gevao SM, Gnirke A, Rambaut A, Garry RF, Khan SH, Sabeti PC. 2014. Genomic surveillance elucidates Ebola virus origin and transmission during the 2014 outbreak. Science 345:1369–1372.

25. Loman NJ, Constantinidou C, Christner M, et al. 2013. A culture-independent sequence-based metagenomics approach to the investigation of an outbreak of shiga-toxigenic escherichia coli o104:h4. JAMA 309:1502–1510.

26. Kundu S, Lockwood J, Depledge DP, Chaudhry Y, Aston A, Rao K, Hartley JC, Goodfellow I, Breuer J. 2013. Next-generation whole genome sequencing identifies the direction of norovirus transmission in linked patients. Clin Infect Dis Off Publ Infect Dis Soc Am 57:407–414.

27. Halachev MR, Chan JZ, Constantinidou CI, Cumley N, Bradley C, Smith-Banks M, Oppenheim B, Pallen MJ. 2014. Genomic epidemiology of a protracted hospital outbreak caused by multidrug-resistant Acinetobacter baumannii in Birmingham, England. Genome Med 6:70.

28. Wilson MR, Naccache SN, Samayoa E, Biagtan M, Bashir H, Yu G, Salamat SM, Somasekar S, Federman S, Miller S, Sokolic R, Garabedian E, Candotti F, Buckley RH, Reed KD, Meyer TL, Seroogy CM, Galloway R, Henderson SL, Gern JE, DeRisi JL, Chiu CY. 2014. Actionable Diagnosis of Neuroleptospirosis by Next-Generation Sequencing. N Engl J Med 370:2408–2417.

29. Greninger AL, Naccache SN, Federman S, Yu G, Mbala P, Bres V, Stryke D, Bouquet J, Somasekar S, Linnen JM, Dodd R, Mulembakani P, Schneider BS, Muyembe-Tamfum J-J, Stramer SL, Chiu CY. 2015. Rapid metagenomic identification of viral pathogens in clinical samples by real-time nanopore sequencing analysis. Genome Med 7:99.

30. Thomson E, Ip CLC, Badhan A, Christiansen MT, Adamson W, Ansari MA, Bibby D, Breuer J, Brown A, Bowden R, Bryant J, Bonsall D, Filipe ADS, Hinds C, Hudson E, Klenerman P, Lythgow K, Mbisa JL, McLauchlan J, Myers R, Piazza P, Roy S, Trebes A, Vattipally SB, Witteveldt J, Consortium S-H, Barnes E, Simmonds P. 2016. Comparison of next generation sequencing technologies for the comprehensive assessment of full-length hepatitis C viral genomes. J Clin Microbiol JCM. 00330–16.

31. Naccache SN, Thézé J, Sardi SI, Somasekar S, Greninger AL, Bandeira AC, Campos GS, Tauro LB, Faria NR, Pybus OG, Chiu CY. 2016. Distinct Zika Virus Lineage in Salvador, Bahia, Brazil. Emerg Infect Dis 22.

